# Adaptive Eye Movement Behavior for Actions in Younger and Older Adults

**DOI:** 10.1101/2025.09.06.674432

**Authors:** Anna Schroeger, Leonard Gerharz, Dimitris Voudouris

**Affiliations:** Department for General Psychology, Justus Liebig University Giessen, Germany; Center for Mind, Brain and Behavior, Universities of Marburg, Giessen, and Darmstadt, Germany

**Author notes:** corresponding author: Anna Schroeger, Alter-Steinbacher Weg 38, 35394 Giessen, Germany.

**Keywords:** gaze, eye movements, aging, interception, prediction

## Abstract

Humans typically combine eye and head movements to visually track and interact with moving objects. If the object’s trajectory can be predicted, humans can shift gaze to the anticipated object’s future location. For instance, when hitting a moving ball, people predictively look at the ball’s bouncing location, allowing them to overcome sensorimotor delays that would otherwise limit visual information uptake from that position. Such delays become pronounced with healthy aging, which might drive a behavioral shift towards stronger reliance on predictive gaze allocation to future positions of interest during visuomotor tasks. To investigate this, we examined gaze and hand movements of older and younger adults who played an iPad-based version of the game ‘Pong’ against an automated opponent. Participants moved a paddle of two different sizes with their finger to intercept a ball moving in two speeds and bouncing off the side walls. We hypothesized that aging would lead to earlier gaze shifts to positions of interest, such as to the final wall bounce location before the ball moved toward the participants’ paddle, and to their own paddle around the moment of interception. As expected, both age groups intercepted fewer balls when using smaller paddles. Contrary to our hypothesis, older participants looked later than younger adults to the final wall bounce. However, they shifted their gaze earlier to their own paddle before interception, especially at the beginning of the experiment. This suggests that aging enhances predictive gaze shifts related to the immediate action. While gaze allocation patterns largely overlap between older and younger adults, aging specifically leads to adaptive eye movements just before interacting with the environment.

## Introduction

Vision plays a central role when performing various tasks, from executing daily activities (Land & Hayhoe, 2001; Wilkie & Wann, 2003) to doing sports (Diaz et al., 2013; Mann et al., 2013). Humans typically look at positions of interest before moving their body to those positions (Coats et al., 2016; Johansson et al., 2001), with such proactive gaze shifts presumably allowing people to sample high resolution visual information related to the body movement (e.g., Hayhoe et al., 2012). Obtaining such relevant visual information as early as possible will in turn foster planning and guidance of the related body movement. For instance, people look at a glass before reaching to pick it up (Johansson et al., 2001; Voudouris et al., 2018) and at their upcoming foothold before stepping on it with high precision (Hollands & Marple-Horvat, 2001; Matthis et al., 2018). When interacting with moving objects, such as when catching a ball, humans combine visual information about the object’s motion with prior information about the underlying dynamics and track the moving object with head and eye movements (de la Malla et al., 2019; Fooken et al., 2016; Spering et al., 2011), especially when the object moves along a rather predictable path. When the moving object follows less predictable paths, prediction of the object’s next position may not be reliable and therefore gaze may not be shifted to the exact position where the target will appear. The resulting mismatches between gaze and target position are corrected by catch-up saccades (Nachmani et al., 2020). Evidently, when sensory feedback processing is delayed, or when the catch-up saccade is executed inaccurately or slowly, retinal mismatches between gaze and target can accumulate and can cause a cascade of inaccuracies that can impair performance (de la Malla & Goettker, 2023; Goettker et al., 2018).

Pursuing a target can be beneficial as it can enhance predictions about the target’s future location (Spering et al., 2011) and improve the timing of interception (Brenner & Smeets, 2011; Diaz et al., 2009). People are still able to intercept a moving object, even if they track the object only briefly (Kreyenmeier et al., 2017; López-Moliner & Brenner, 2016), because this brief glimpse provides enough visual information to predict the ball’s motion and its final destination (Fooken & Spering, 2019). Tracking a moving object only briefly can also allow gaze shifts to other positions of interest. For instance, when performing bat-and-ball interceptive actions, people predictively shift their gaze around the future position of the ball’s bounce, sometimes even at a position along the trajectory that the ball will follow after the bounce (Diaz et al., 2013; Hayhoe et al., 2012). These predictive gaze shifts occur even earlier when the ball moves with higher speeds (Diaz et al., 2013), likely because visual information about the ball’s bounce and rebound is available for shorter durations, and it is important to sample this information as it can facilitate tracking the ball after the rebound. Indeed, people can pursue a bouncing ball after its rebound with latencies of just around 50 ms (Hayhoe et al., 2012), highlighting that the gaze shifts to the ball’s position after the rebound is based on predictions through prior knowledge and sampled visual information from the ball’s trajectory. These flexible gaze shifts presumably allow to overcome inherent sensorimotor delays that can otherwise impair human visuomotor behavior.

Delays in processing incoming sensory input and in executing motor outputs are an inherent characteristic of the sensorimotor system. Such delays become more pronounced during healthy aging (Hultsch et al., 2002; Seidler et al., 2010) and are the combined result of degradation in various sensory, motor and cognitive functions. Indeed, aging typically comes with poorer sensory processing (Klever et al., 2019; Owsley, 2011, 2013), impaired musculoskeletal functionality (Doherty et al., 1993), increased motor variability (Seidler et al., 2010) and slower motor performance (Kanekar & Aruin, 2014). That older adults typically perform their actions slower can be explained by such physiological alterations, but these do not exclude the possibility that sensorimotor slowing may be an actively chosen compensation strategy to maintain higher accuracy of the executed movement. Studies using fMRI (Biehl et al., 2017) and fNIRS (Ward et al., 2018) revealed that older adults have increased neural activity in visual brain areas during motion perception tasks, which might indicate compensation by involving additional neural substrates to work out the motion dynamics. Another compensatory mechanism that older adults could rely on when confronted with pronounced sensorimotor delays that may affect actions requiring prompt and timely behavior, is to exploit predictive strategies and obtain sensory information ahead of time. By doing so, they may gain more time to plan and perform the respective body movements. However, to this day, there is no consensus as to whether aging leads to a stronger reliance on predictive control (Coats & Wann, 2012; Kanekar & Aruin, 2014; Vandevoorde & Orban De Xivry, 2019). Specifically, when there is no urge for prompt visuomotor behavior, older adults do not appear to rely on predictive gaze shifts (Gerharz et al., 2024). However, when performing sequential arm movements or walking, older adults shift their gaze to their next target before their ongoing movement is over (Chapman & Hollands, 2006; Coats et al., 2016). Although these suggest stronger reliance on predictive gaze shifts, this proactive gaze allocation may also be a byproduct of older adults simply moving their body slower than younger adults. We also recently demonstrated that older adults employ predictive gaze shifts to positions where high spatiotemporal precision is required (Gerharz & Voudouris, 2025). Specifically, when participants had to hit a moving target at a pre-determined hit area, older adults shifted their gaze to that area much earlier than younger adults, highlighting that aging can cause an adaptive shift in predictive gaze allocation when performing tasks that require high spatiotemporal precision.

In this study, we are interested in whether aging is characterized by stronger reliance on predictive gaze shifts during a continuous, gamified task that varies in complexity. Healthy young and older participants played an iPad version of the game ‘Pong’, where they had to move with their index finger a virtual paddle to intercept a ball that was moving between their own paddle and that of an automated opponent. The ball was returned from the opponent’s paddle in various angles that resulted in collisions with the side walls at various positions and eventually approached the participant’s paddle in various directions. In separate blocks of trials, we set the paddle to be either small or large, and the ball to move either at a slow or fast speed, both of which allowed us to manipulate the complexity of the task. We expect poorer interception rates with smaller paddles and faster balls. We also expect poorer interception rates in older than younger adults. More central to our aims, we hypothesize that participants look at relevant positions before the ball arrives at those positions. Specifically, based on previous findings (Diaz et al., 2013; Mann et al., 2013), we expect participants to predictively look at around the final wall bounce, which will allow them to sample visual information about the ball’s rebound direction and promptly pursue it as it moves toward their own paddle. Likewise, we expect participants to look around their own paddle before the ball arrives there, as has been shown in other interception tasks (de la Malla et al., 2019; Gerharz & Voudouris, 2025). We expect these predictive gaze shifts to occur earlier with balls that move faster (i.e., Diaz et al., 2013; main effect of speed), as the time during which the ball is at those relevant positions will be shorter with faster balls, and this can make it more important to look at those positions well before the relevant information arises there. Our main interest is in whether aging influences predictive gaze allocation, and whether this is sensitive to the underlying complexity. If aging leads to stronger reliance on predictive gaze shifts to positions of interest, we expect older adults to look at those positions earlier than younger participants (main effect of age), specifically under more challenging conditions (interaction between age and ball speed or paddle size).

**Figure 1.**
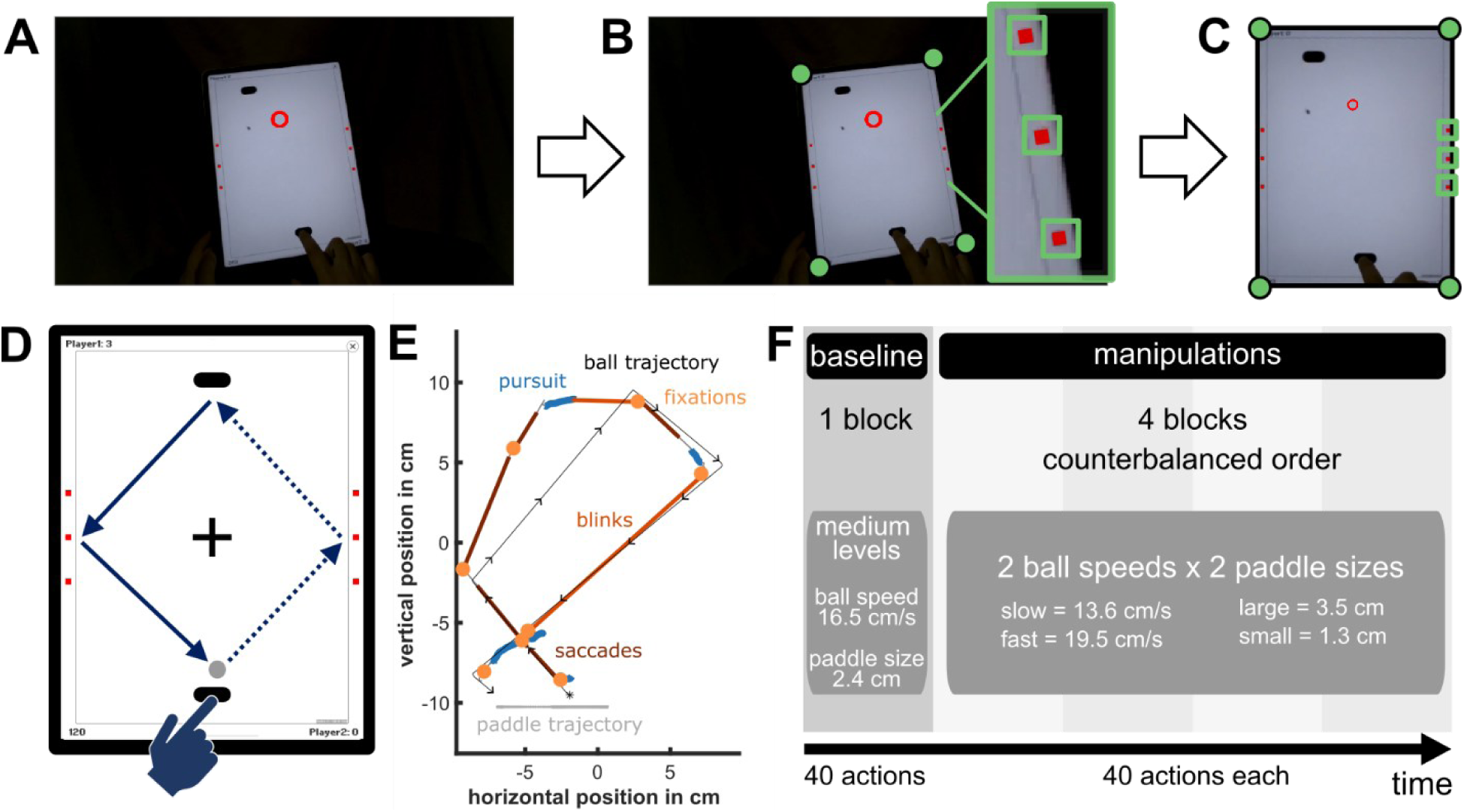
Overview of the video game ‘Pong’ and the methods. ***A)*** Illustrative frame captured by the front camera of the mobile eye tracker. ***B)*** In each video frame, we detected either the three reference rectangles on each side or the four corners of the iPad. ***C)*** The detected reference locations were used to transform gaze coordinates and video input onto the iPad using Computer Vision Algorithms. ***D)*** One action cycle was defined as the time between two actions of the participant, with the opponents’ action in between. During a single action cycle, several wall bounces could occur. An action cycle breaks down into a receding phase (dotted lines), when the ball moves from the participant’s paddle towards the opponent, and an approaching phase (solid lines), when it moves back to the participant’s paddle. ***E)*** Illustrative action cycle. The ball followed the trajectory (black line) as it started moving from the paddle (*), bounced once at the left-side wall, before bouncing back from the opponent and once at each wall on the way back. The participant tracked the ball with a combination of saccades (dark red parts) and smooth pursuit eye movements (blue parts) and fixated (orange circles) on the bounce locations and other locations on the ball’s trajectory. Blinks (red parts) were typically combined with large gaze shifts (like saccades). ***F)*** Illustration of the experimental design. All participants started the experiment with a baseline block of medium ball speed and paddle size levels. The order of the four experimental blocks (each ball speed combined with each paddle size) was counterbalanced across participants.

## Results

### Interception performance

First, we tested whether the manipulations of paddle size and ball speed affected interception performance measured as the relative frequency of hits across all actions. Performance was poorer in older than younger adults, *F*(1, 36) = 10.35, *p* = .003, η_p_^2^ = .22. It was also poorer when the task was more demanding, as there were fewer hits when balls moved faster than slower, *F*(1, 36) = 130.39, *p* < .001, η_p_^2^ = .78, and when paddles were smaller than larger, *F*(1, 36) = 226.26, *p* < .001, η_p_^2^ = .86. Importantly, the detrimental effects of fast balls and small paddles were larger for older than younger adults, reflected in interactions between ball speed and age group, *F*(1, 36) = 8.60, *p* = .006, η ^2^ = .19, and between paddle size and age group, *F*(1, 36) = 11.79, *p* < .002, η ^2^ = .25, and the respective post-hoc comparisons (*p* = .006 and *p* = .001). Additionally, fast balls specifically affected performance for small paddles and only lesser so, when paddle size was large, as indicated by an interaction between ball speed and paddle size, *F*(1, 36) = 47.31, *p* < .001, η ^2^ = .57. Post-hoc comparisons confirmed that the difference between fast and slow balls was larger for small compared to large paddles (*p* < .001).

**Figure 2.**
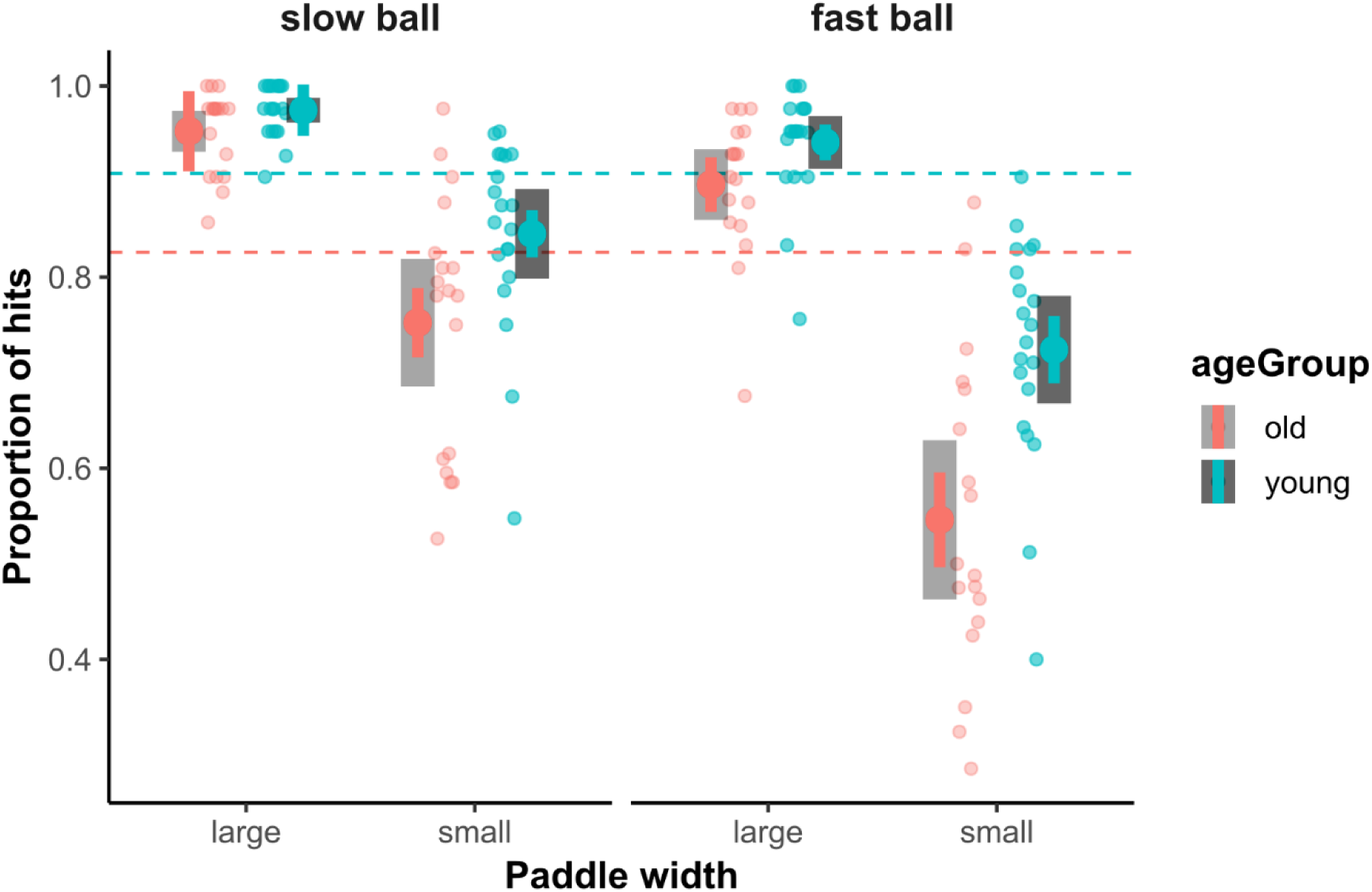
Interception performance across conditions. Dotted lines indicate baseline performance (intermediate ball speed and paddle size), which already reveals age differences in performance. Colored error bars indicate within-subject 95% confidence intervals, grey-scale error bars indicate between-subject confidence intervals.

### Description of eye and hand movements

Similar to a previous study using the same design (Schroeger et al., 2024), our participants adjusted their eye and hand movement behavior throughout the action cycle. They specifically increased the time of pursuing the ball in the second (approaching phase) compared to the first half (receding phase) of the action cycle, peaking shortly before their own action (interception; Fig. 3A). Pursuit of the opponent’s paddle (Fig. 3B) or their own paddle (Fig. 3C) was less frequent and restricted to times around the related actions. The rate of fixations was relatively constant across the duration of the action cycle (Fig. 3D), while the rate of saccades was largely reduced around the time of the participants’ or the opponent’s interception (Fig. 3E). Participants strategically increased their blinks shortly after their own interception. This is not surprising because missing visual information from that moment has likely the least interference with task performance, and participants kept blink-frequency at a minimum while the ball was approaching their paddle (Fig. 3F). Gaze allocation as described by the above-mentioned parameters appears rather similar between younger and older adults, although older adults seem to pursue the ball more frequently as this approaches the opponent’s paddle (Fig. 3A). Older adults moved the paddle with their hands later than younger adults did (Fig. 3E). A cross-correlation between the time-curves of older and younger adults revealed a delay of 5% relative time (*r* = .97) which corresponds to ∼160 ms based on an average action cycle duration of 3.19 s.

**Figure 3.**
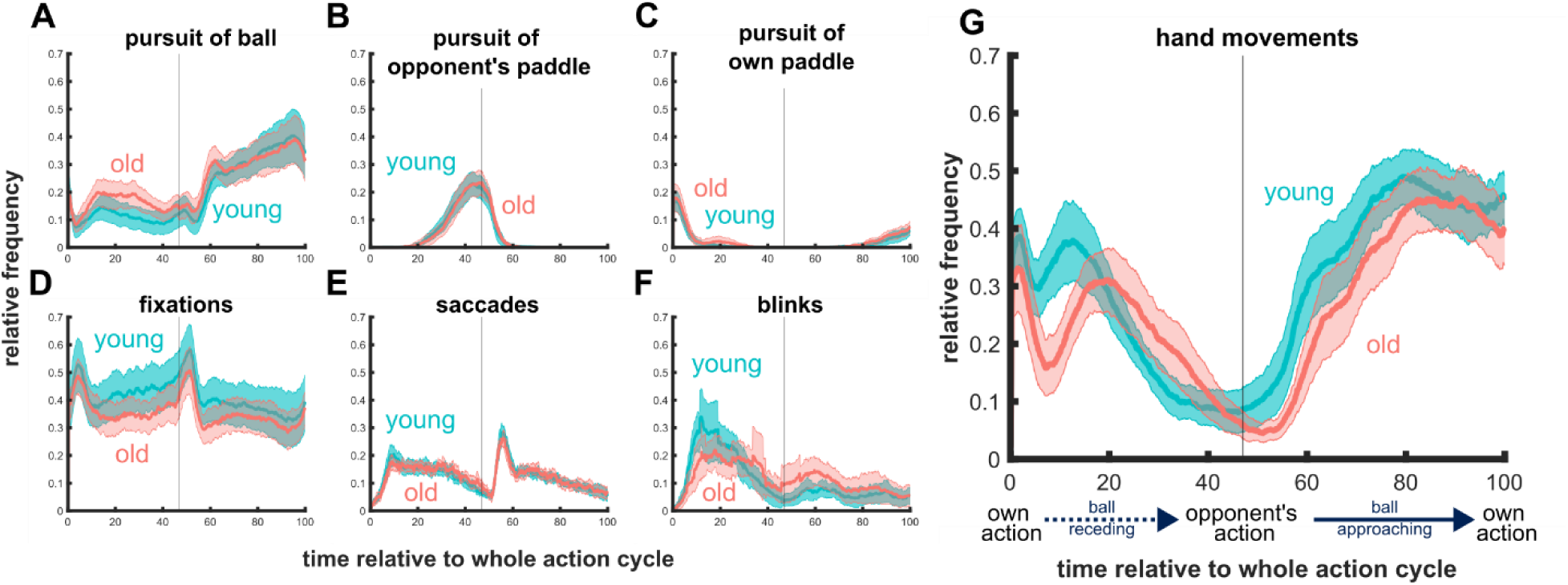
Distribution of gaze- and hand-related events over time course of an action cycle. ***A)*** Participants pursued the ball mainly when it was approaching them, ***B)*** they pursued their opponent’s paddle around the time of the opponent’s action, and ***C)*** pursued their own paddle only rarely after they intercepted themselves. Additionally, participants’ ***D)*** fixation rate did not fluctuate much but it dropped after the opponent returned the ball to the participant. ***E)*** Participants reduced their saccade rate around the moments of interception, and ***F)*** they blinked mainly during the receding ball phase. While the overall movement patterns of older adults resemble those of the younger adults, the peak of blinks in the receding phase are attenuated and fixations decrease in frequency across the whole action cycle. Older adults seem to pursue the ball already more frequently in the receding phase, whilst younger adults initially only rarely pursue but show a more pronounced increase in the approaching phase. ***G)*** Most prominently, older adults move their hands later than younger adults, and even the whole distribution appears shifted in time. In between the start (0% of action cycle) and end frames (100% of action cycle), the ball bounced once from the opponent, which is marked by a grey vertical line at 47%, since this was the average bounce time across all action cycles.

### Predictive eye movements

We were mainly interested in whether participants would perform predictive gaze shifts to positions of interest in order to obtain ahead-of-time visual information that could be relevant for their action. To this end, we looked at when participants directed their gaze i) to the wall where the ball bounced for the last time before moving toward the participants’ paddle, and ii) to their own paddle around the time of interception. We expressed these times relative to when the ball touched the wall and the paddle, respectively. We used two repeated measures ANOVAs on the time of the gaze shift to these two locations, including age group as between- and paddle width and ball speed as within-subjects factors. We report oculomotor prediction, indicating the time when the eyes first arrived around the bounce location (within ∼2 degrees of visual angle from the position of interest), and we express this time relative to the moment of the bounce^1^, with positive values representing stronger predictive gaze shifts (i.e. earlier gaze at the respective location). As becomes evident in Figure 4A-B, most participants directed their gaze to the final wall and to their own paddle (Fig. 4A) *before* the ball arrived at that location, indicating predictive gaze shifts to locations of interest, final wall bounce: *t*(37) = 13.26, *p* < .001, *d* = 2.15, and interception bounce: *t*(37) = 10.63, *p* < .001, *d* = 1.72. We were mostly interested in whether aging would influence the reliance on predictive gaze shifts in the examined task and whether the predictions depend on the ball speed and paddle width.

**Figure 4.**
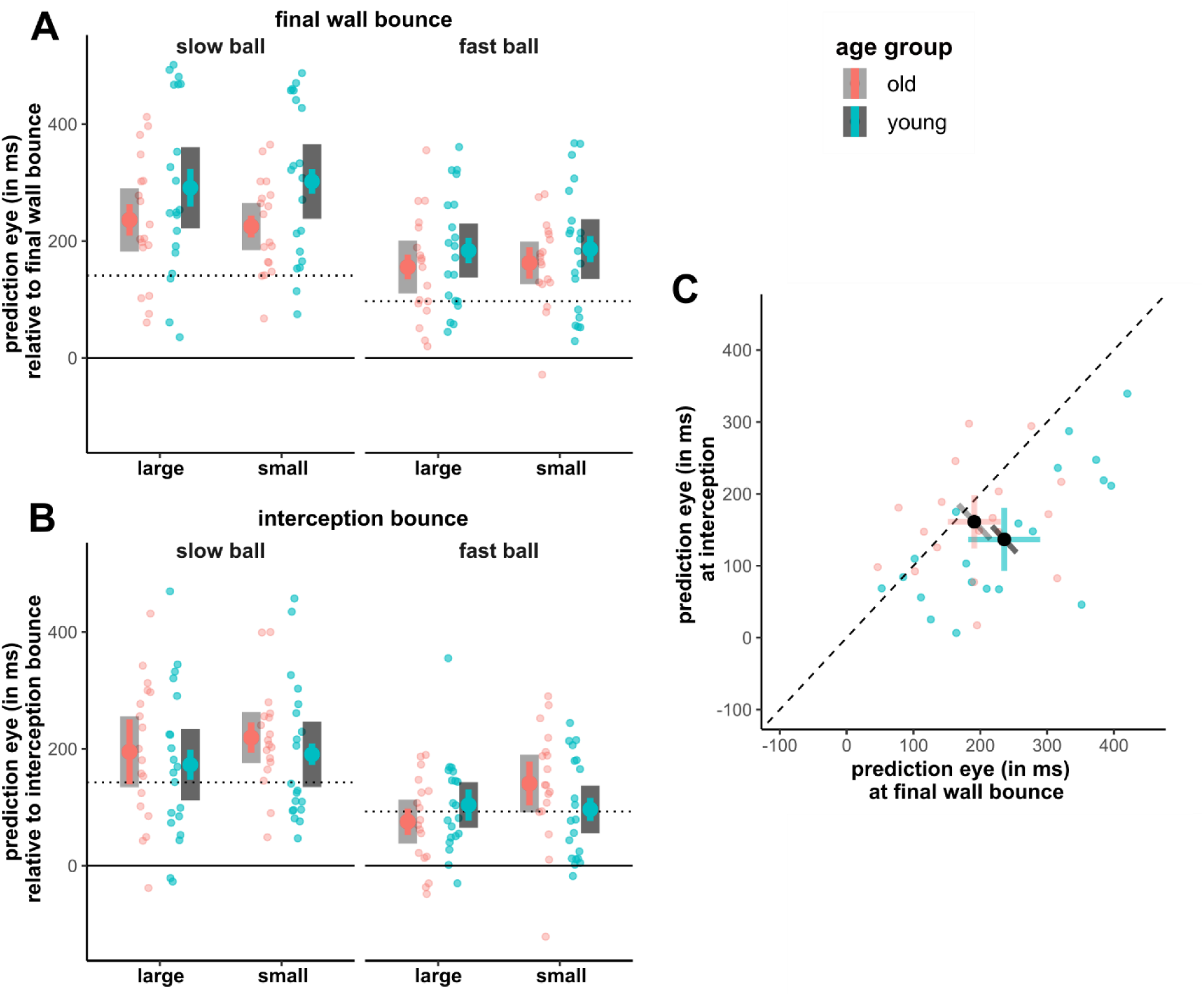
Predictive eye movements. ***A)*** Time of gaze arriving at the final wall bounce area relative to the ball’s arrival at that location per ball speed and paddle size. Positive values indicate prediction; negative values indicate delay. The grey dotted line indicates when the ball arrived at the 2 deg distance from the bounce position, which represents the spatial error of our eye-tracker. That the values are higher than this dotted line further supports the idea that participants make predictive gaze shifts and are not solely pursuing the moving target. ***B)*** Same as A but for the interception bounce (paddle). ***C)*** Pairwise comparisons of gaze arriving at the two locations (aggregated across ball speeds). While younger adults reduce their prediction from the final wall bounce to the interception location (most data points are below the dotted line), older adults do not systematically change their gaze behavior between the two locations of interest. For the last wall bounce younger adults seem to shift their gaze earlier than older adults, while the pattern appears inverted for the interception location. Colored error bars indicate within-subject 95% confidence intervals, grey-scale error bars indicate between-subject confidence intervals.

*Final wall bounce*. The repeated measures ANOVA on predictive gaze shifts at the final wall bounce revealed a main effect of ball speed, *F*(1, 36) = 105.40, *p* < .001, η_p_^2^ = .75. While slow balls elicited predictive gaze shifts on average 264 ms before the ball bounced off the wall, fast balls came with predictive gaze shifts that occurred relatively later, on average 172 ms before the ball bounced (Fig 4A). This effect was further elucidated by a significant interaction between age group and ball speed, *F*(1, 36) = 5.07, *p* = .030, η_p_^2^ = .12. Post-hoc contrasts revealed that younger adults largely scaled their predictive gaze shifts according to the ball’s speed: their gaze shift to the wall bounce location was on average 112 ms earlier when the ball moved slower than faster (*p* < .001). Older adults also scaled the timing of their predictive gaze shift, but more moderately, as they looked at the wall bounce on average 71 ms earlier when the ball moved slower than faster (*p* < .001). Yet, the effect of ball speed on predictive gaze allocation was systematically more pronounced in younger than older adults (*p* < .031). Overall, these results are against our hypotheses and previous findings (e.g., Diaz et al., 2013), because they show earlier gaze shifts around the position of interest with *slower* ball movements. One possible reason for this finding could be that we define the gaze shift as any gaze within 2 deg from the position of interest (wall or paddle). Tracking the ball closely along its trajectory to that position can yield earlier gaze shifts when the ball moves slower than faster, because the ball reaches the 2 deg area with more time left until reaching the exact bounce position. However, when we re-ran our analysis by only looking at fixations within the same area, instead of *any gaze,* the pattern of results remains qualitatively similar. Please note that the dotted lines in Figure 4 A and B indicate when the ball reaches the 2 degrees area of interest. This illustrates that gaze was on average *before* the ball.

*Interception bounce.* Also, for the interception bounce, results revealed a significant main effect of ball speed, *F*(1, 36) = 68.55, *p* < .001, η ^2^ = .66, with earlier predictive gaze shifts when the ball moved slower (194 ms) than faster (104 ms, Fig. 4B). Additionally, there was a significant main effect of paddle width, *F*(1, 36) = 4.92, *p* = .033, η ^2^ = .12. Participants looked at their paddle ∼137 ms before interception when the paddle was large but ∼162ms before interception when the paddle was small.

*Exploratory analysis on bounce location across age groups.* Given that the timing of predictive gaze shifts appeared to change between the two bounce locations, we additionally ran an exploratory repeated measures ANOVA on the timing of these predictive gaze shifts across bounce locations and age groups. Indeed, predictive gaze was shifts occurred earlier when the ball was travelling toward the last wall bounce (∼213 ms) compared to the interception location (∼149 ms), as evident in a significant main effect, *F*(1, 36) = 20.46, *p* < .001, η_p_^2^ = .36. Although we did not observe a main effect of aging, the analysis revealed a significant interaction between bounce location (wall vs. interception) and age group, *F*(1, 36) = 5.91, *p* = .020, η_p_^2^ = .14. Post-hoc analyses revealed that younger adults shifted their gaze 99 ms earlier to the final wall bounce compared to the interception location (*p* < .001), indicating that younger adults relied less on prediction for their interception (and potentially used more pursuit eye movements). Meanwhile, for older adults the latency of the eye movement to the wall did not differ significantly from that to the interception location (*p* = .158, see Fig. 4C).

### Relationship between interception performance and gaze allocation

The previous results raise the question whether predictive gaze shifts to positions of interest provide any functional advantage and possibly benefit performance. To test this, we ran a between-subject correlation between (a) the average time of gaze shift to the positions of interest, and (b) the proportion of hits, after collapsing all the data across all paddle width and ball speed conditions. Earlier oculomotor predictions to the final wall bounce significantly improved participants’ performance, *r*(36) = .54, *p* < .001. Figure 5A shows the linear fits split up for each condition with similar trends in all four conditions and for both age groups. However, the timing of the predictive gaze shifts to the interception location (paddle) was not correlated with performance scores, *r*(36) = .03, *p* = .843 (Fig. 5B).

**Figure 5.**
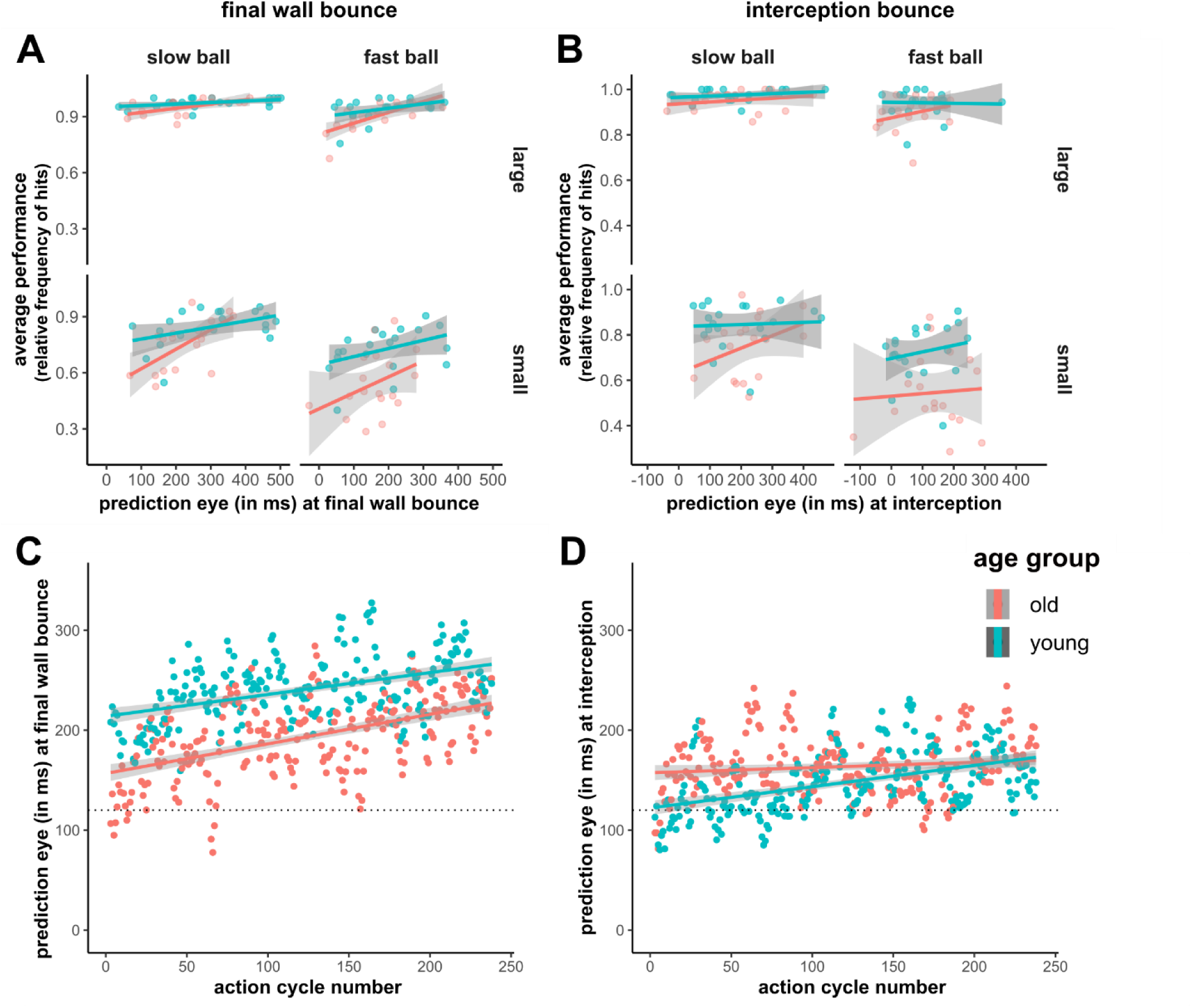
Predictive gaze allocation across conditions and over time of the experiment. **(A, B)** Correlation between predictive gaze shifts and interception performance for each experimental condition. **(C, D)** Changes in predictive gaze allocation over the duration of the experiment**. A)** In all conditions, participants with earlier predictive gaze shifts to the final wall bounce performed better in the game. Please note that only one overall correlation, aggregated across conditions, was run because performance was too close to ceiling in the conditions with the large paddle. **B)** Predictive gaze allocation to the interception location did not correlate with performance. **C)** Predictive gaze allocation to the final wall bounce is evident from the beginning of the experiment, but predictive gaze shifts became more pronounced with more exposure to the task, for both groups. **D)** Predictive gaze shifts to the interception location (paddle) were evident from the beginning of the experiment, especially in older adults. However, the modulation over the time course of the experiment is smaller in size. The grey dotted lines in C and D indicate the moment when the ball arrived at the 2 deg distance from the bounce location (serving as a stricter criterion for “prediction”). Please note that the data in panels C and D were filtered using a moving average with a window of 5 trials.

We additionally tested whether the predictive gaze shifts change over the time of the experiment by running a linear regression with action cycle number and age group explaining the prediction of the final wall bounce (see Fig. 5C). We found a significant main effect of action cycle, indicating that both age groups shifted their gaze earlier to the wall bounce location over the time of the experiment, β = 0.28, *t*(475) = 6.17, *p* < .001. There was also a main effect of age group, with younger adults demonstrating predictive gaze shifts of ∼47 ms shorter latencies than older adults, β = 47.06, *t*(475) = 7.41, *p* < .001. However, the analysis revealed no interaction (*p* = .391) indicating that the learning rate was similar between both groups: according to the linear fit, with each action cycle participants predicted approximately 0.3 ms more than in the previous one.

A second regression tested whether participants relied stronger on predictive gaze shifts to the interception location (paddle) with more exposure in the task. This revealed a similar but smaller main effect of action cycle number, β = 0.13, *t*(476) = 3.27, *p* = .001. Over the time of the experiment participants only slightly improved their interception prediction at a rate of 0.13 ms per action cycle. In contrast to the final wall bounce, older adults now demonstrated predictive gaze shifts of ∼16 ms shorter latencies than those of younger adults, which lead to a significant main effect of age group, β = −15.51, *t*(476) = −2.71, *p* = .007. It is striking from the figure below that older adults adopt a predictive gaze strategy from the beginning of the experiment, whereas younger adults develop such strategy as they are exposed more in the task. This suggests that older adults rely more on predictive gaze shifts to positions of interest from the early phases of the task, whereas younger adults seem to initially adopt a different strategy, which they abandon as they perform more trials. Eventually, the behavior of younger adults aligns with the behavior that older adults adopted from early on. Despite this qualitative observation, the analysis revealed no significant interaction (*p* = .083).

## Discussion

We tested whether older adults rely more strongly on predictive gaze allocation, which could serve as a compensation for the increasing visuomotor processing delays that manifest during aging. To do so we compared interception performance and gaze allocation between a group of older (55-69 years) and a group of younger adults (20-35 years). We used a gamified interception task of different levels of complexity. Our results revealed that aging led to poorer interception rates and stronger consequences of the increased task complexity. In line with the assumption that sensorimotor delays increase with age (Hultsch et al., 2002; Seidler et al., 2010), decrements in interception performance of older adults were likely caused by their delayed hand movements when controlling the paddle that they used to intercept the moving target. Importantly, both older and younger adults shifted gaze predictively to bounce locations, specifically the locations of the final wall bounce and of their theoretical interception. Although the average timing of the predictive gaze shifts was similar between age groups, we observed differences throughout an action cycle and the course of the experiment. For instance, both age groups looked at the final wall bounce in a predictive manner, but this appeared to be more pronounced in younger adults (see Fig. 5C). The predictive gaze shifts to that location became more pronounced with more exposure to the task, while the rate of change was similar between age groups. This indicates that both younger and older participants learned the game’s underlying dynamics and tailored their gaze behavior accordingly. However, older adults relied more strongly on predictive gaze shifts to the interception location than younger adults did at the beginning of the experiment (see Fig. 5D). With more exposure to the task, participants of both age groups used even earlier predictive gaze shifts. Thus, for positions where interactions are more important, older adults appear to adopt predictive gaze strategies from early on, while younger participants evolve such strategies with more exposure to the task. If older adults deployed compensatory processes these might rather be focused on the moment of interception. During this final part of the ball trajectory both age groups engage largely in pursuit eye movements, but older adults appear to additionally maintain predictive gaze shifts. Importantly, earlier gaze shifts to the final wall bounce were related to better interception rates, whilst earlier gaze shifts to the interception location (paddle) did not appear related to interception performance.

### Performance differences

Our results clearly showed that interception performance is impaired in older adults, a finding that might be attributed to increasing sensorimotor delays and the poorer sensory processing that is often associated with healthy aging (Hultsch et al., 2002; Seidler et al., 2010). We further showed that older adults suffered more from difficult task conditions, such as when the ball moved at faster speeds and when the interception paddle was smaller. Although previous work has also shown a reduced number of hits during interception in older than younger adults (de Dieuleveult et al., 2019), such age-related decline in manual performance is not always reported (Gerharz & Voudouris, 2025). One possible reason for this may be that the older participants in the latter study employed predictive gaze strategies that were more pronounced than those of younger participants. In other words, predictive gaze shifts may allow older adults to preserve interception performance. Even if older adults employed predictive gaze shifts in our study, these gaze shifts had similar latencies to those of younger adults and might have not been predictive enough to provide the necessary visual information well in-advance and keep performance at similar levels as reached by younger adults.

### Movement patterns across an action

Overall, older and younger adults used similar gaze allocation strategies to fulfill the task demands, but older adults appeared specifically delayed in their hand movements when controlling the paddle. Both age groups limited gaze behaviors that deteriorate the visual input, like blinks and saccades, to less critical moments in time, for instance when the ball was still far away from their next action. This is important because visual information uptake is compromised during blinks and saccades, and therefore it can be crucial to reduce this behavior around moments when critical visual events can happen (Hoppe et al., 2018; Hoppe & Rothkopf, 2016; Schroeger et al., 2024; Shalom et al., 2011). Indeed, a closer look into the distribution of blinks in the current study suggests that the strategic pattern of increasing blinks occurs shortly after participants returned the ball back to the opponent and before the opponent intercepts. During this period any changes in the ball’s and opponent’s trajectory is relatively uninformative for the participant’s next action.

Both older and younger adults only occasionally pursued the ball when it approached the opponent but largely increased pursuit when it approached their own paddle. These pursuit frequencies peak shortly before the moment of interception. Participants might have strategically increased pursuit at this critical time to improve their predictions about the ball’s future trajectory which could facilitate the accuracy and timing of their interception (Brenner & Smeets, 2011; Schroeger et al., 2024; Spering et al., 2011). Our results indicate that both younger and older adults make use of this strategy. Still, the difference between the first and second half of the action cycle appears slightly more distinct in younger adults (see Fig. 3). Throughout the whole action cycle older adults spent less time fixating. Overall, eye movement patterns are comparable between age groups, with potentially slightly more strategic timing for younger adults.

The most prominent difference between age groups emerged in the hand movement frequencies. While the overall pattern is comparable, it appears that older adults move the paddle with their finger with approximately 160 ms of a delay. Similarly, Gerharz et al. (2024) revealed longer hand movement latencies in older adults when reaching to hit visual targets. In a two-segment aiming task, Rand & Stelmach (2011) showed that older adults exhibit longer hand movement, acceleration and deceleration times than younger adults. All these findings might indicate a degradation in various sensory, visual, motor and cognitive functions (Hultsch et al., 2002; Klever et al., 2019; Owsley, 2011; Seidler et al., 2010) or a compensation strategy of slowing movements to keep accuracy of movements high.

### Predictive gaze shifts across age groups

We hypothesized that older adults would rely more strongly on predictive gaze shifts, possibly to compensate for the sensorimotor delays. Although average behavior did not reveal any evidence for this hypothesis, we found an interaction between age group and location of interest and we observed different patterns of predictive gaze allocation between age groups throughout the experimental procedure. Specifically, younger participants employed earlier predictive gaze shifts to the final wall bounce location than to their own paddle. The fact that predictive gaze shifts to the paddle were initiated relatively later than those to the final wall might be related to participants increasing the amount of pursuit shortly before interception (see Fig. 3). However, older adults showed no such difference between the timing of their predictive gaze shifts despite them still increasing pursuit frequency during the final approach phase of the ball. This is probably because older adults adopted predictive gaze allocation patterns –both to the final wall bounce and to their own paddle– from the early phases of the experiment, whereas younger participants did so only for the final wall bounce location (see Figure 5C-D). Although we did not find direct evidence of a general increase in predictive gaze shifts in older adults, Figure 5D reveals that the two age groups adopted different gaze strategies throughout the task. During the early phases of the experiment, older adults adopted clearly stronger predictive gaze allocation patterns than younger adults for their own interception. This is also suggested by a trend for an interaction between number of action cycles and age group. As participants performed more trials, they increased the reliance on predictive gaze shifts to the paddle just before interception, but this increase appears steeper in younger than older adults. Thus, when exposed with unknown dynamics, older adults seem to employ predictive gaze shifts to locations of interest whereas younger adults initially rely on such a strategy to a lesser extent. However, after exposure to the dynamics, the differences between the age groups resonate. This is striking because it shows that younger adults are slower in adopting gaze strategies that are already obtained by the older participants in earlier stages.

Both age groups learned the ball’s dynamics and tailored their gaze behavior accordingly. While older adults initiate predictive gaze shifts toward the final wall bounce later relative to younger adults, the learning rate as indicated by the change in the latency of the gaze shifts is similar between younger and older adults. Comparable learning rates were previously observed in for low-level saccadic adaptation processes (Huang et al., 2017) and predictive gaze strategies (Gerharz et al., 2024), demonstrating preserved capacity for oculomotor improvement across younger and older adults. However, for the interception location, the learning rate tended to be smaller for older than younger adults. This presumably arises because the two age groups started at different levels in their timing of predictive gaze allocation, and there probably is a threshold beyond which it is unnecessary to use even earlier predictive gaze shifts, for instance because one needs to also pursue the moving target for some time. The earlier predictive gaze shifts toward the interception location of the older adults might have already started at an elevated level compared to those of younger adults because of an inherent change in gaze allocation strategy to account for the longer sensorimotor delays in aging. Other potential explanations might be related to difficulties that older adults have with pursuing moving objects. Previous research revealed that older adults exhibit decreased smooth pursuit gain, indicating that they are less able to keep their gaze aligned with the target (Moschner & Baloh, 1994; Sharpe & Sylvester, 1978). Additionally, they need more time to initiate pursuit as indicated by increased pursuit latencies (Sharpe & Sylvester, 1978). Accordingly, older adults might put less trust in their pursuit eye movements and maintain predictive strategies to compensate for reduced pursuit gain.

We found that ball speed and occasionally paddle size impacted how early participants shifted their gaze to the bounce area. In contrast to our predictions, both age groups shifted their gaze to the positions of interest earlier with slower than faster balls. This makes sense if participants do not change their spatial predictions. To predict earlier for fast balls, participants would need to make larger gaze shifts, or in other words predict the trajectory spatially further. Only for the interception bounce, we additionally found an effect of paddle size. Participants predict the interception location earlier when they control a small compared to a large paddle. This finding is in line with the idea that participants have to adapt to increased task demands. When the paddle is small the potential impact surface is smaller, and interception has to be even more accurate. It might therefore be advantageous to shift gaze earlier to the predicted interception location because this leaves more time to potentially correct these predictions or adapt the hand movement.

## Conclusion

Older adults employ similar movement strategies but seem delayed in their hand movements during interception. In contrast to younger adults, they maintain predictive gaze shifts until the moment of acting, whilst younger adults instead reduced their predictions shortly before acting and appeared to largely rely on pursuit eye movements. In general, prediction of the final wall bounce was associated with better game performance and was increased over the time of the experiment. Importantly, both groups learn to increase their predictions at the same rate.

## Methods

### Participants

A total of 38 participants completed the experiment. We contrasted a group of older adults (N = 18, age range: 55-69 years, mean age = 62.2 years, sd = 4.8 years, 7 male) with a group of younger adults (N= 20, age range: 20-35 years, mean age = 26.1 years, sd = 3.7 years, 3 male). Older adults revealed slightly worse visual acuity (mean_old_ = 0.24 logMAR, sd_old_ = 0.31; mean_young_ = −0.15, logMAR, sd_young_ = 0.14) and contrast sensitivity (mean_old_ = 1.59 logCS, sd_old_ = 0.39; mean_young_ = 1.90, logCS, sd_young_ = 0.14), assessed with the Freiburg Visual Acuity Test (Bach, 1996, 2024). One older participant was excluded because the video recording suggests that he moved the glasses during a break and data quality dropped afterwards. Younger participants were recruited from university student pools and were compensated either with 8€/hour or course-credits for their participation. Older adults were community-dwelling and were recruited from university mailing lists and personal contacts of the authors. They were compensated with 8€/hour. Participants gave informed consent prior to participating in the study. This experiment is part of a project approved by the ethics committee of the University of Giessen (approval number: LEK-2021-0028) and was conducted in accordance with the Declaration of Helsinki (2013; except for §35, pre-registration).

### Materials

The Pong game was played on an iPad 12 Pro (2732 x 2048 px, 264 ppi) and was similar to the game used in a previous study (Schroeger et al., 2024). A grey dot representing the ball (radius: 0.19 cm = 20 px) moved between two black rectangular paddles with rounded edges, presented on a uniform white background (screen size: 26.3 cm x 19.7 cm, see Fig. 1C). The paddles could move only in the horizontal direction. The movement of the one paddle was controlled by the participant while the other paddle (‘opponent’) was automated. The opponent was programmed to move whenever the ball approached that paddle. It followed the horizontal position of the ball including a small variable temporal delay and variable speed, and it almost always intercepted the ball directing it toward the side walls and the participant’s paddle. Every 7 actions (see below), the horizontal position of the opponent’s paddle was slightly shifted with respect to that of the ball, making the opponent miss the ball. The paddles had a fixed height of 100 px (∼1 cm) and in separate blocks we set the widths of both paddles to be either *small* (140 px) or *large* (360 px), which corresponded to 1.3 and 3.5 cm, respectively (see Fig. 1F). The ball moved linearly with a constant speed between the two paddles and bounced back from the side walls. In separate blocks, the speed of the ball was either *slow* (23.6 px/frame) or *fast* (33.8 px/frame), which corresponded to ∼13.6 cm/s and ∼19.5 cm/s, respectively. An initial block served as baseline and practice using an intermediate paddle size of 250 px (2.4 cm) and ball speed of 28.6 px/frame (16.5 cm/s). The distance between participants’ eyes and the iPad ranged between ∼50-60cm, which means that 1 cm on the iPad corresponds to ∼0.96-1.15 degrees of visual angle. To achieve variable ball trajectories the bounce angle from the opponent was chosen randomly in each interception as a random number between 35 and 55 degrees relative to perpendicular (vertical axis). The emergent angle equaled the incidence angle for all wall bounces. That implies that all bounces except for the bounce from the opponent’s paddle were predictable.

We recorded eye and head movements using the Tobii Glasses Pro 3 and analyzed the data in MATLAB (The MathWorks Inc., 2022) using the GlassesViewer (Niehorster et al., 2020) and self-written code and R version 4.1.2 (R Core Team, 2021) using RStudio (RStudio Team, 2016). Gaze data was sampled at 100 Hz, the scene at 25 Hz, and the IMU at ∼100 Hz. Participants could move their paddle by sliding their index fingertip along the horizontal direction of the iPad. All stimuli were displayed on the iPad, which also served to save paddle and ball positions at 60 Hz. All data was interpolated to 100 Hz prior to any further processing (e.g. applying filters).

### Procedure

All participants filled out a questionnaire on their demographic information (age, gender) and completed a computerized test of visual acuity and contrast sensitivity using Landolt-C shapes from the Freiburg Vision Test (Bach, 1996, 2024). Older adults were additionally tested for their cognitive functions using the Montreal Cognitive assessment and all passed with a score of at least 26 (Nasreddine et al., 2005). A short calibration and validation of the eye tracker was performed prior to the experiment (GlassesValidator, Niehorster et al., 2023). In the Pong game, instead of separate trials, participants performed a series of actions until one of the players missed the ball, which prompted the game to restart. We later analyze each action taken by the participant and therefore divide the series of actions into action cycles, defined as the time from the first data point after an action of the participant (i.e. interception of ball) to the data point of their next action (i.e. next ball interception after the opponent returned the ball; see Fig. 1C). Participants completed 5 experimental blocks resulting from the 2 (ball speed) x 2 (paddle size) experimental design and the additional preceding practice block, in which the ball speed and paddle size were in-between those used in the experimental conditions (see Fig. 1F). The baseline block was always performed first in the sequence, while the subsequent four blocks were presented in a random order to counterbalance across participants. Each block required participants to perform a minimum of 40 actions irrespective of whether it was a successful interception or a miss. A block then ended with the next miss (either by the participant or the opponent) and was followed by a short break which participants could extend based on their needs. Each series of actions started with a fixation cross at the center of the screen, presented for 2.5 to 3 ms drawn from a uniform distribution. A ball then appeared at the center of the iPad and immediately started moving towards the opponent’s side, bouncing back from the walls. Once the opponent intercepted the ball, it continued to the participant’s paddle, again bouncing off walls. The game was continuous until one of the two players missed the ball, and the next series of actions started with the presentation of a fixation cross at the center of the tablet. The validation of the eye-tracker was repeated after the end of the experiment. The whole session took around 1-1.5h.

### Eye and Head Movement Analysis

Once per participant we manually temporally aligned the data recorded by the eye tracker with the data recorded by the iPad using the scene video and a green rectangle that was briefly displayed on the screen at the beginning of the game. We converted eye coordinates into gaze position on the iPad in MATLAB using object detection and homography transformation with the functions *bwboundaries*, *fitgeotrans* (projective), *transformPointsForward*, *imwarp* and a color mask based on HSV color space, all implemented in MATLAB. To do so, within each frame of the scene camera we detected six red rectangles positioned at the iPads’ frame using a color mask and the function *bwboundaries*. If this detection failed, we detected the iPad’s corners using *bwboundaries* (see Fig. 1 A-C). Using these 6 (red rectangles) or 4 (iPad’s corners) reference locations, we mapped the gaze data onto the iPad’s screen with the functions *fitgeotrans* (projective), and *transformPointsForward*, and transformed the video frame to its original resolution with *imwarp*.

We labeled saccades and blinks based on the original eye movements (prior to gaze mapping) as implemented in the GlassesViewer (Niehorster et al., 2020) choosing the adaptive velocity algorithm from Hessels at al. (2020). Given typical saccade durations (Gandhi & Katnani, 2011; Leigh & Zee, 2015) we changed labels of saccades that lasted equal to or longer than 200 ms to blinks. We classified pursuit eye movements based on the mapped gaze data: Initially we selected all intervals with gaze velocities larger than 7 cm/s and of at least 100 ms duration which corresponds to approximately half the velocity of the slowest target. Gaze and velocity traces were filtered using a 3^rd^ order Savitzky-Golay filter with a length of 11 frames (110 ms). Of these we only included intervals in which the gaze moved in the same direction (same angle +/− 30 degrees) as one of the three objects (ball and two paddles). Finally, we associated each pursuit interval with its corresponding pursuit target (ball, own paddle, opponent’s paddle) by selecting the target with the most similar movement direction. For example, when the ball is close to one of the paddles, ball and paddle might move in very similar directions, but the ball never moves perfectly horizontal – in contrast to the paddles. If the pursuit was more closely aligned to horizontal motion, we associated it with the paddle movement, and otherwise with the ball. Remaining intervals in the gaze traces of at least 100 ms duration, which were neither labeled as saccades, blinks or pursuit, nor missing data, indicated nearly stationary gaze positions and were therefore labeled as fixations. All intervals shorter than 100 ms or with missing data were not labeled (16% of gaze traces).

Hand movements were determined based on the Savitzky-Golay-filtered speed of the participant’s paddle (3^rd^ order, length of 11 frames), as this was extracted from the positional data provided by the iPad. A hand movement was based on the paddle speed being larger than 5 cm/s for at least 100 ms, irrespective of the direction of motion.

### Statistical Analysis

We first segregated the data streams into separate action cycles, with the start frame being the moment after the ball bounced from the participants’ paddle and the end frame being the moment of the next bounce from the participant’s paddle. In between the start (0% of action cycle) and end frames (100% of action cycle), the ball bounced once from the opponent, which was set to be at the 47%-mark of a single action cycle, since this reflects the ratio of the average durations of the receding and approaching phase. Please note that these phases can differ in duration since they depend on the angle in which the ball bounces from both paddles. We then excluded all action cycles where more than 45% of the gaze data was missing – either between two consecutive interceptions, or, in case of the first interception attempt, between the moment when the ball left the center of the iPad and the next interception of the participant – or when technical issues occurred (1.5% of action cycles).

We analyzed participants’ performance by calculating the relative frequency of successful interceptions for each condition relative to all action cycles in this condition. To test for differences between age groups and conditions, we ran a mixed-design ANOVA, with ball speed and paddle size as within-, and age group as between-subject factors and Holm-corrected post hoc tests based on the marginal means.

To identify different strategies in their movement patterns, we descriptively compared eye and hand movement frequencies over the time of an action cycle. For this purpose, we calculated the relative frequency of each type of movement for each moment (0.1% steps) within the action cycle. To test for delays, we compared hand movement traces of older and younger adults by calculating a cross-correlation: we correlated intervals of 201 values (20% time) and shifted the older adults’ interval by −10 to 20% relative time in steps of 1% compared to the younger adults’ trace, to test which shift revealed the highest correlation. We repeated this analysis for 20 intervals (ranging from 20% relative time to 58% relative time as central time point in the 201 values window), each shifted by 2% of relative time, and then aggregated the correlation coefficients (per delay) across these 20 repetitions by taking the median. In other words, for each of the 31 temporal shifts between groups (−10 to 20%), we aggregated correlation coefficients across 20 different temporal segments of the hand movement trace. We then identified the delay revealing the highest correlation.

To analyze whether participants used predictive gaze strategies, whether age groups differed in the amount of prediction, and whether interception performance correlates with prediction, we picked two potential indicators of prediction (see also, Diaz et al., 2013): the time when the gaze reached the ball’s final wall bounce location, and the time when gaze reached the theoretical interception location irrespective of whether they hit or missed the ball (both relative to the ball reaching these locations). Given that the spatial accuracy and precision of the eye-tracker is limited, we defined a region of ∼2 degrees of visual angle (∼2 cm) around the actual locations. Another reason for this approximate area was that previous research has shown that participants rather do not exactly direct their gaze at the bounce location but rather an ‘elevated’ location that facilitates tracking after the bounce (Diaz et al., 2013). We subtracted the ball’s bounce-time from the arrival time of gaze within each region of interest (positive values indicating prediction) and aggregated the prediction times for each participant and the two locations. We ran two mixed-design ANOVAs for the two bounce locations each with the between-subject factor age group and the two within-subject factors ball speed and paddle size. We additionally ran a mixed-design ANOVA with the between-subject factor age group, and the within-subject factor location and two post-hoc contrasts on the difference between locations in each age group to evaluate the interaction. To test whether predictive gaze shifts benefit performance, we aggregated the prediction times and the performance per participant and tested an association by correlating them.

To analyze the change in predictive gaze over the time of the experiment we aggregated gaze timing across participants per action cycle. We ran a linear regression on each measure of predictive gaze, with age group and action cycle number as predictors. For illustration, we additionally used a moving average of size 5 to smoothen the data.

For statistical analyses we used R version 4.1.2 (R Core Team, 2021) and RStudio (RStudio Team, 2016) and interpret significance relative to an alpha value of .05. We report effect sizes as partial eta squared. We additionally used the following packages: R.matlab (Bengtsson et al., 2018), readxl (Wickham & Bryan, 2025), Rmisc (Hope, o. J.), dplyr (Wickham et al., 2025), ez (Lawrence, 2016), emmeans (Lenth, 2025), rstatix (Kassambara, 2023), ggplot2 (Wickham, 2016).

## Acknowledgements

We thank Jelmer de Vries for programming the first version of the game, and Viktoria Wild and Marie Mengel for assistance with data collection.

## Data availability statement

Data and Materials will be made public on osf upon acceptance for publication.

## Declaration of generative AI and AI-assisted technologies in the writing process

During the preparation of this work the author(s) used ChatGPT and Perplexity.ai in order to improve readability of the manuscript. After using this tool/service, the author(s) reviewed and edited the content as needed and take(s) full responsibility for the content of the publication.

## Footnotes

1 Given that the overall accuracy of the eye tracker might lead to errors above 2 deg of visual angle, we additionally corrected the data in this specific analysis based on a validation procedure at the beginning and the end of the experiment, because we rely on spatial values. We present new versions of Figures 4 and 5 with the corrected data in the Supplement as Figures S1 and S2.

