## Supplemental Figures for "Adaptive Eye Movement Behavior for Actions in Younger and Older Adults"

* corresponding author

Given that the overall accuracy of the eye tracker is limited, we additionally corrected the data and Figures on predictive gaze shifts to bounce locations because these rely on spatial values. We therefore shifted the position based on the average accuracy of two 9-point validation procedures (Niehorster et al., 2023) conducted prior and after the experimental task. Visual examination of the 'corrected' Figures indicates consistency with those presented in the main manuscript.

### Predictive eye movements

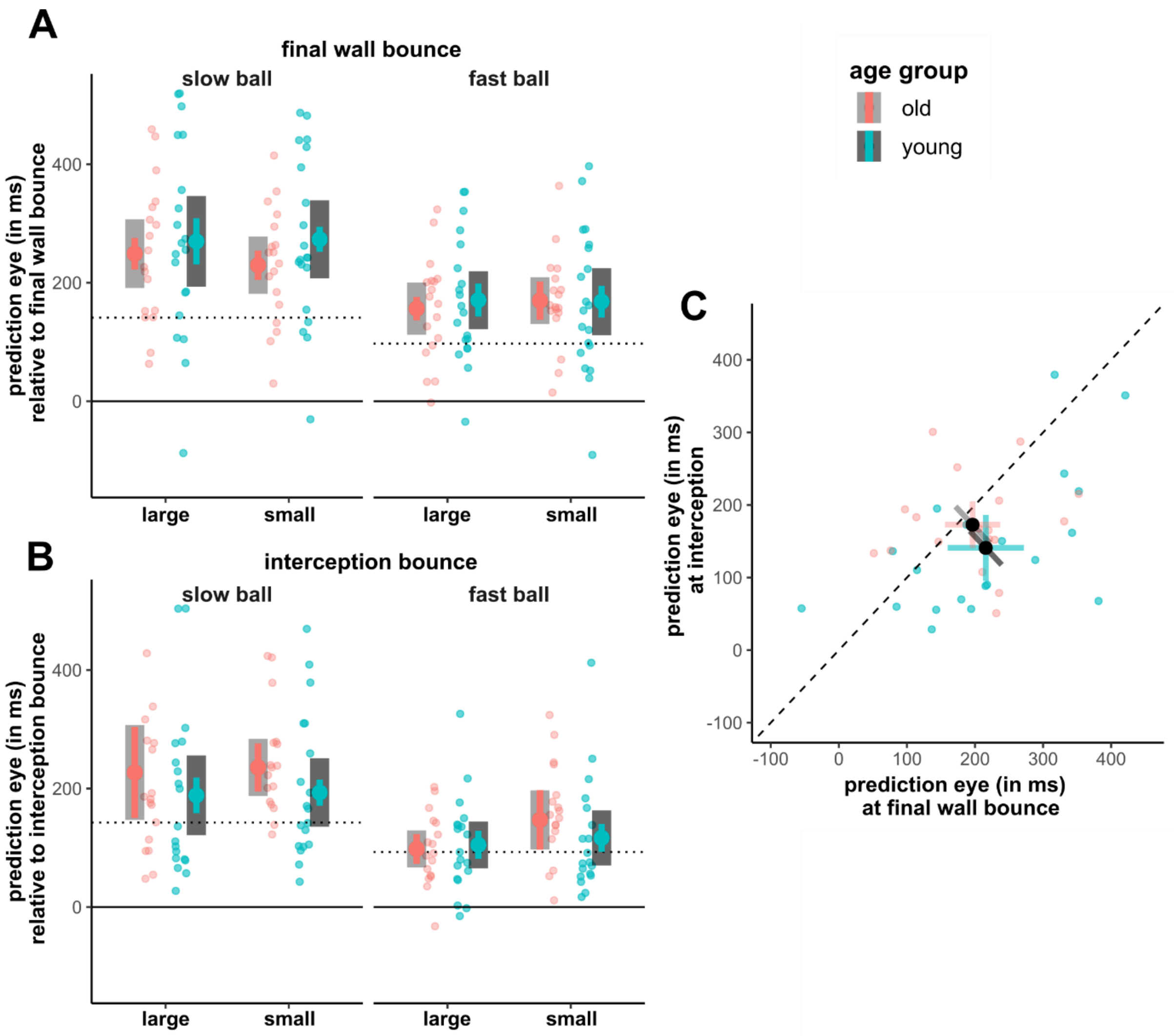


***Figure S1. Predictive eye movements for 'corrected' data. A)*** *Time of gaze arriving at the final wall bounce area relative to the ball's arrival at that location per ball speed and paddle size. Positive values indicate prediction; negative values indicate delay. The grey dotted line indicates when the ball arrived at the 2 deg distance from the bounce position, which represents the spatial error of our eye-tracker.* ***B)*** *Same as A but for the interception bounce (paddle).* ***C)*** *Pairwise comparisons of gaze arriving at the two locations (aggregated across ball speeds). While younger adults reduce their prediction from the final wall bounce to the interception location (most data points are below the dotted line), older adults do not systematically change their gaze behavior between the two locations of interest. For the last wall bounce younger adults seem to shift their gaze earlier than older adults, while the pattern appears inverted for the interception location. Colored error bars indicate within-subject 95% confidence intervals, grey-scale error bars indicate between-subject confidence intervals.*

### Relationship between interception performance and gaze allocation

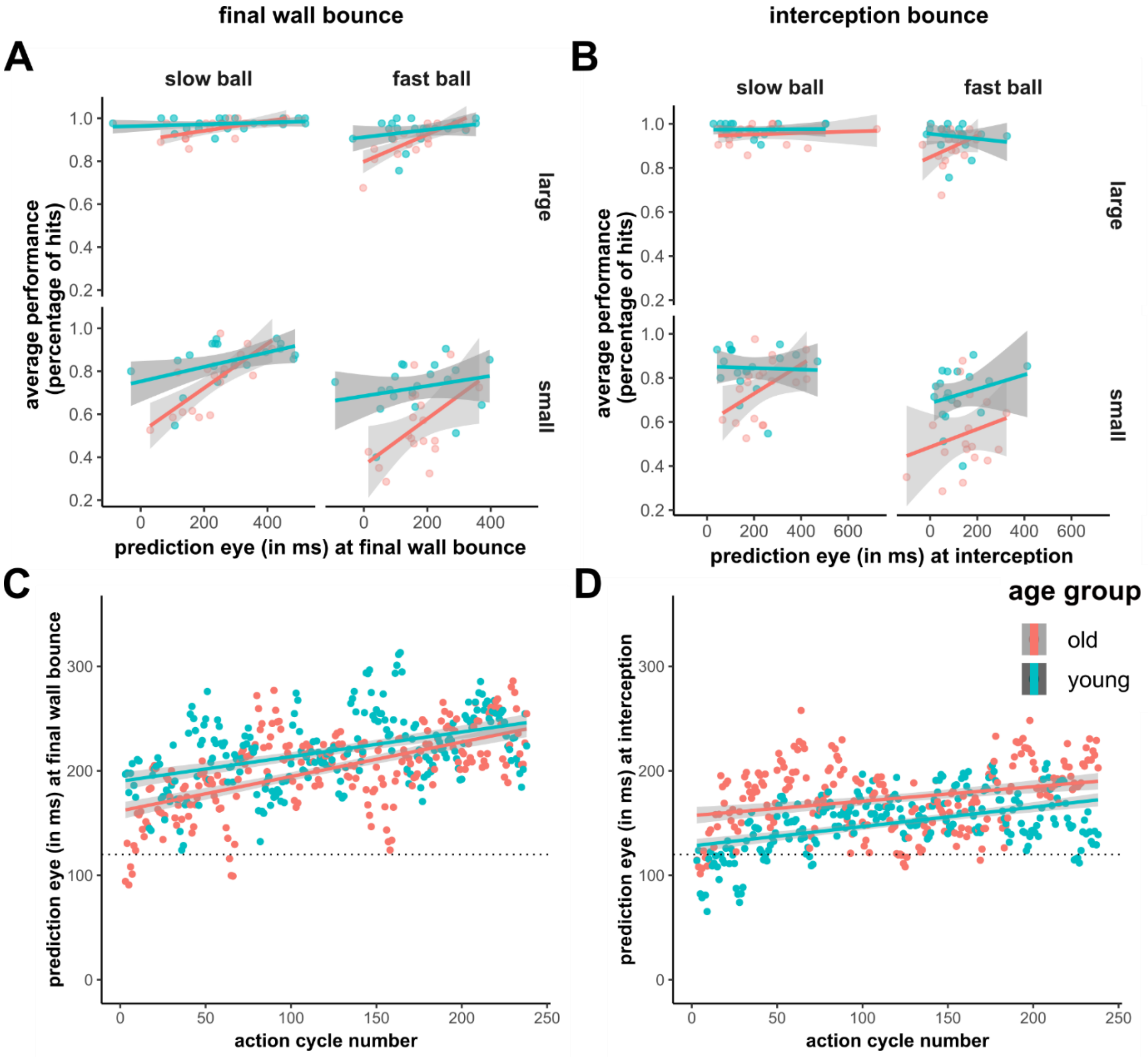


*__Figure S2. Predictive gaze allocation across conditions and over time of the experiment for 'corrected' data__. (A, B) Correlation between predictive gaze shifts and interception performance for each experimental condition. __(C, D)__ Changes in predictive gaze allocation over the duration of the experiment. __A__) In all conditions, participants with earlier predictive gaze shifts to the final wall bounce performed better in the game. Please note that only one overall correlation, aggregated across conditions, was run because performance was too close to ceiling in the conditions with the large paddle. __B)__ Predictive gaze allocation to the interception location did not correlate with performance. __C)__ Predictive gaze allocation to the final wall bounce is evident from the beginning of the experiment, but predictive gaze shifts became more pronounced with more exposure to the task, for both groups. __D)__ Predictive gaze shifts to the interception location (paddle) were evident from the beginning of the experiment, especially in older adults. However, the modulation over the time course of the experiment is smaller in size. The grey dotted lines in C and D indicate the moment when the ball arrived at the 2 deg distance from the bounce location (serving as a stricter criterion for "prediction"). Please note that the data was filtered using a moving average for plots C and D.*

#### *References*


Niehorster, D. C., Hessels, R. S., Benjamins, J. S., Nyström, M., & Hooge, I. T. C. (2023). GlassesValidator: A data quality tool for eye tracking glasses. *Behavior Research Methods*, *56*(3), 1476–1484. https://doi.org/10.3758/s13428-023-02105-5
